# What drives phenotypic divergence among coral clonemates?

**DOI:** 10.1101/514430

**Authors:** Iliana B Baums, Meghann K Devlin-Durante, Dana W Williams, Dustin Kemp

## Abstract

Evolutionary rescue of populations depends on their ability to produce phenotypic variation that is heritable and adaptive. DNA mutations are the best understood mechanisms to create phenotypic variation, but other, less well-studied mechanisms exist. Marine benthic foundation species provide opportunities to study these mechanisms because many are dominated by isogenic stands produced through asexual reproduction. For example, Caribbean acroporid corals are long lived and reproduce asexually via breakage of branches. Fragmentation is often the dominant mode of local population maintenance. Thus, large genets with many ramets (colonies) are common. Here, we observed phenotypic variation in stress response within genets following the coral bleaching events in 2014-and 2015 caused by high water temperatures. This was not due to genetic variation in their symbiotic dinoflagellates (*Symbiodinium ‘fitti’*) because each genet of this coral species typically harbors a single strain of *S. ‘fitti’*. Characterization of the microbiome via 16S tag sequencing did not provide evidence for a central role of microbiome variation in determining bleaching response. Instead, epigenetic changes were significantly correlated with the host’s genetic background, the position of the sampled polyps within the colonies (e.g. branch versus base of colony), and differences in the colonies’ condition during the bleaching event. We conclude that microenvironmental differences in growing conditions led to long-term changes in the way the ramets methylated their genomes contributing to, but not fully explaining, the differential bleaching response. This research provides novel data to understanding intra-genet variability in stress phenotypes of sessile marine species.

## Introduction

Acclimatization is a non-genetic process by which an individual can increase its stress tolerance after exposure to a stressor, such as temperature anomalies (see Palumbi *et al.* 2014 for a coral example). Acclimatization can lead to phenotypic variability in stress response among clonemates. Non-mutation based mechanisms resulting in phenotypic variability in isogenic lines include stochastic gene expression, errors in protein synthesis, protein promiscuity and epigenetic modifications (reviewed in Payne & Wagner 2019). Protein synthesis, for example, has rates of mistranslation far exceeding those of DNA point mutations (Yanagida *et al.* 2015). Of these, only epigenetic mutations have been studied as a mechanism for acclimatization in marine foundation fauna (reviewed in Eirin-Lopez & Putnam 2019). Interestingly, non-mutation based changes to phenotypes can prolong survival of a genet until such phenotypes become permanent (Yanagida *et al.* 2015), however, these processes are not well understood. Somatic mutations may also change phenotypes among ramets of the same genet and have become a focus for studies on marine foundation fauna (Devlin-Durante *et al.* 2016; Reusch & Boström 2011; Santelices *et al.* in press; Van Oppen *et al.* 2011) although no reports have yet tied a change in phenotype to a somatic mutation in marine foundation species. Understanding of all mechanisms that produce phenotypic variability is essential to estimate the evolvability of threatened marine species (Payne & Wagner 2019).

The large-scale bleaching event during 2014-2015 within the Florida Keys provided an unprecedented opportunity to understand the role of acclimatization and phenotypic variability in structural foundation species of shallow Caribbean coral reefs. Because reef-building corals harbor intracellular symbionts (family Symbiodiniaceae), discerning the relative contribution of host and symbiont to holobiont acclimatization can be difficult. However, the Caribbean elkhorn coral, *Acropora palmata*, has an uncomplicated symbiosis: it associates with just one symbiont species (*Symbiodinium ‘fitti’*) and most colonies (ca. 70%) also harbor only one strain of *S. ‘fitti’* over space and time (Baums *et al.* 2014) -a finding predicted by theory (Parkinson & Baums 2014) and one that has been observed for other cnidarian-*Symbiodinium* associations (e.g. Andras *et al.* 2011; Thornhill *et al.* 2009). Thus, *A. palmata* is a good model to discern the coral host’s acclimatization response and phenotypic variability to heat stress.

Like many reef-building corals, *A. palmata* frequently reproduces via fragmentation (an asexual process), sometimes forming large, monoclonal stands (Baums *et al.* 2006; Foster *et al.* 2007; Pinzón *et al.* 2012; Williams *et al.* 2014). These colonies represent iterations of the same host-symbiont combination (i.e. they are replicates), experiencing similar environmental conditions. Initial surveys of *Acropora palmata,* during both summer 2014 and 2015 bleaching events in the Florida Keys documented a range of bleaching responses. This response varied between reefs but also within single, monoclonal stands of *A. palmata* (Fig 2). Thus, coral clonemates were observed to exhibit different bleaching susceptibilities despite data showing that they share identical (clonal) *S. fitti* symbiont communities, begging the question as to what mechanisms account for such phenotypic variability. The answer may inform our understanding of how reefs might survive climate change and has implications when choosing genets for restoration. Thus, we set out to explore whether differences in the microbiome other than dinoflagellates (e.g. prokaryotes), and/or micro-environmental differences, such as shading or exposure to water movement, induced epigenetic changes in the host genome that could explain differences in bleaching susceptibilities among ramets of the same genet.

DNA methylation occurs at the cytosine bases of eukaryotic DNA, which are converted to 5-methylcytosine by DNA methyltransferase (DNMT) enzymes. DNA methylation is the most highly studied mechanism in epigenetics and is often used to elucidate how a phenotype is modified without altering the genetic code. Additional epigenetic changes include histone modifications, chromatin remodeling and gene regulatory mechanisms involving small noncoding RNAs (Danchin *et al.* 2011). Epigenetic modifications can rapidly produce new phenotypes in response to a change in the environment without mutations in the underlying genetic sequence (Finnegan 2002; Richards 2008).

There is limited data on methylation mechanisms in invertebrates. Current evidence has shown that invertebrate genomes are far less methylated than vertebrates (Gavery & Roberts 2010, 2013; Lyko *et al.* 2010; Olson & Roberts 2014; Rivière 2014; Suzuki *et al.* 2007). Additionally, DNA methylation is predominately found in gene bodies in which highly expressed housekeeping genes are hypermethylated and regulated and/or inducible genes are hypomethylated (Dimond & Roberts 2016; Elango *et al.* 2009; Gavery & Roberts 2010; Hunt *et al.* 2010; Sarda *et al.* 2012). Hypermethylation of those genes essential for biological function is thought to imply that they are “protected” from plasticity in transcriptional opportunities. Such plasticity would be inherently lethal in housekeeping genes (Roberts & Gavery 2012), thereby, gene body methylation in corals is correlated with stable and active transcription (Dixon *et al.* 2017).

In contrast, the inducible gene’s limited methylation may facilitate, albeit passively, specific transcriptional opportunities including increasing sequence mutations, access to alternative transcription start sites, and exon skipping (Roberts & Gavery 2012). There is evidence for a direct relationship between DNA methylation and phenotypic plasticity as seen in the determination of caste structure in both honeybees and ants (Kucharski *et al.* 2008), maternally inherited epigenetic patterns that influence the expression of the *agouti* gene in mice (Wolff *et al.* 1998), and the differential methylation of the human *NR3C1* gene in newborns depending on prenatal maternal mood (Oberlander *et al.* 2008). Scleractinian corals can display strong differences in their DNA methylation in response to stress demonstrating that *de novo* DNA methylation may be a driving mechanism for phenotypic plasticity in acclimation (Putnam *et al.* 2016).

Here, we sampled six different genets from four different reef sites in Florida and sampled each genet six times (either across the same colony 6 times, or different ramets of the same genet for a total of six samples). Samples were taken from the upward facing side of branches or the bases/trunks of colonies six weeks after colonies had experienced a thermal stress event that resulted in differential bleaching. We determined how many *S. fitti* strains were present in each sample and sequenced the 16S gene to characterize variability in the prokaryotic community. We then applied a reduced sequencing technique sensitive to the methylation status of Cytosine called MethylRad (Wang *et al.* 2015) to identify sites that were differentially methylated between reefs, genets, position within the colony, and peak bleaching status.

## Methods

### Sample Collection

During routine monitoring of established *Acropora palmata* study sites (Williams et al 2014) in September 2014 severe bleaching was observed. Colony condition including a ranking of bleaching severity was recorded (Williams et al 2017) and detailed photos were taken of each colony. Bleaching severity ranks ranged from 0 indicating normal colored tissue to 5 for severe bleaching indicating intact tissue that was lacking any color (appeared white). These ranks were applied based on the overall colony bleaching severity of the colony as a whole in the field, ranks were applied by a single observer (D Williams). Bleaching severity was monitored in October and again in November, and routine quarterly monitoring was resumed in 2015 and continued through the 2015 bleaching event.

During November 2014 surveys, multiple tissue samples were collected from several precise locations on identified colonies. The tissue collection locations were selected based on their bleaching condition which varied within and between colonies. Collection sites included four reef sites within about 20km on the Florida reef tract (Fig 1). We sampled six different genets from these four reef sites in Florida and sampled each genet six times (either across the same colony 6 times, or different ramets of the same genet for a total of six samples (Table 1)). Visual assessment of the tissue condition of the sample spot was scored as normal color (NC) or slightly pale (SP), very pale (VP) or Bleached (BL). The location of the sample within the colony was classifies as either exposed branch (high irradiance) or non-branch (base, lower irradiance). Tissue was collected using a small chisel or snippers and the tissue sample was immediately preserved underwater (30 seconds after collection) using RNAlater (Sigma Aldrich).

**Figure 1.**
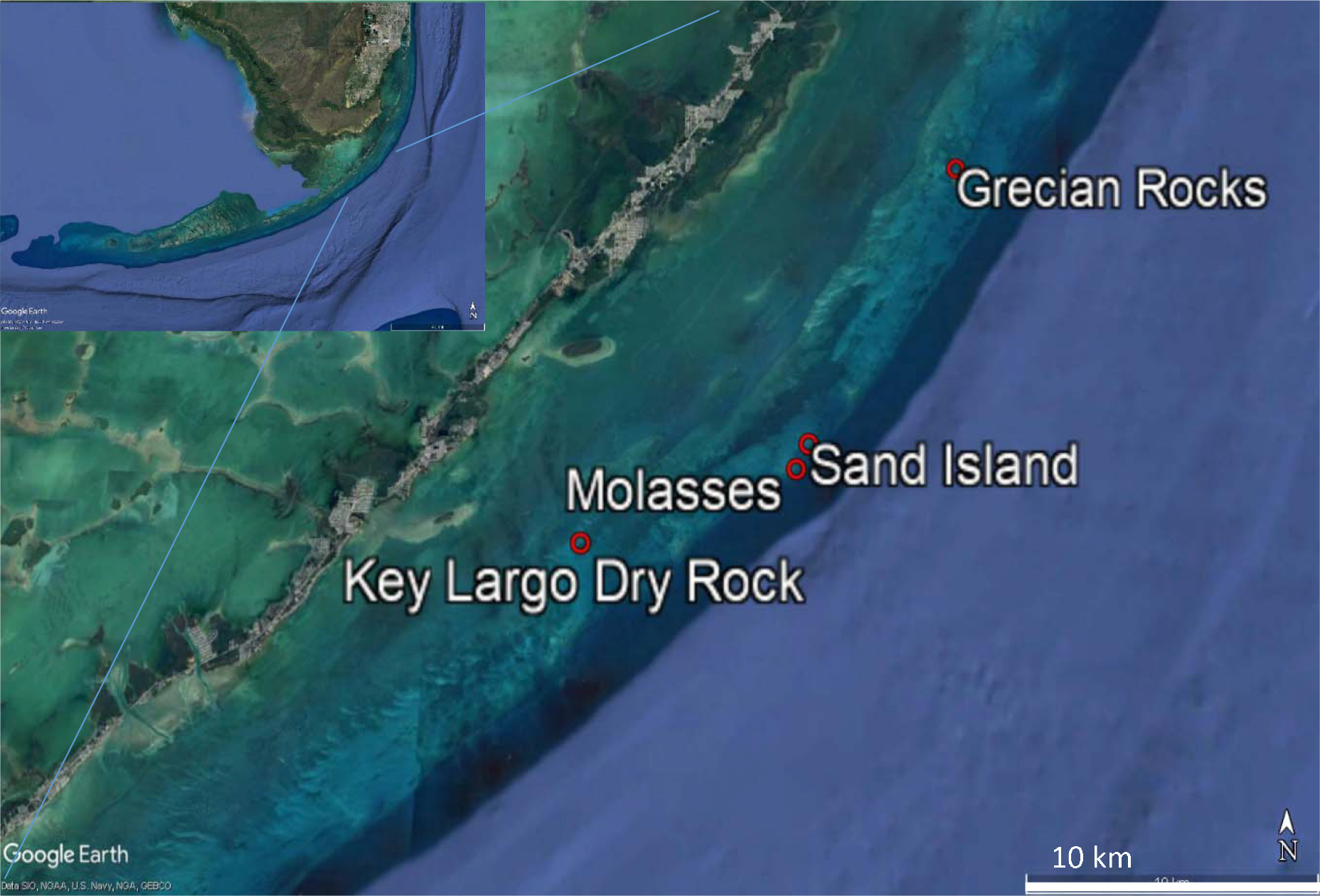
*Acropora palmata* samples were obtained from four reef sites within the Florida Reef tract, Grecian Rocks, Sand Island, Molasses and Key Largo Dry Rocks. Distance between Grecian Rocks and Key Largo Dry Rocks is about 20 km. Maps created in Google Earth Pro.

**Table 1.**
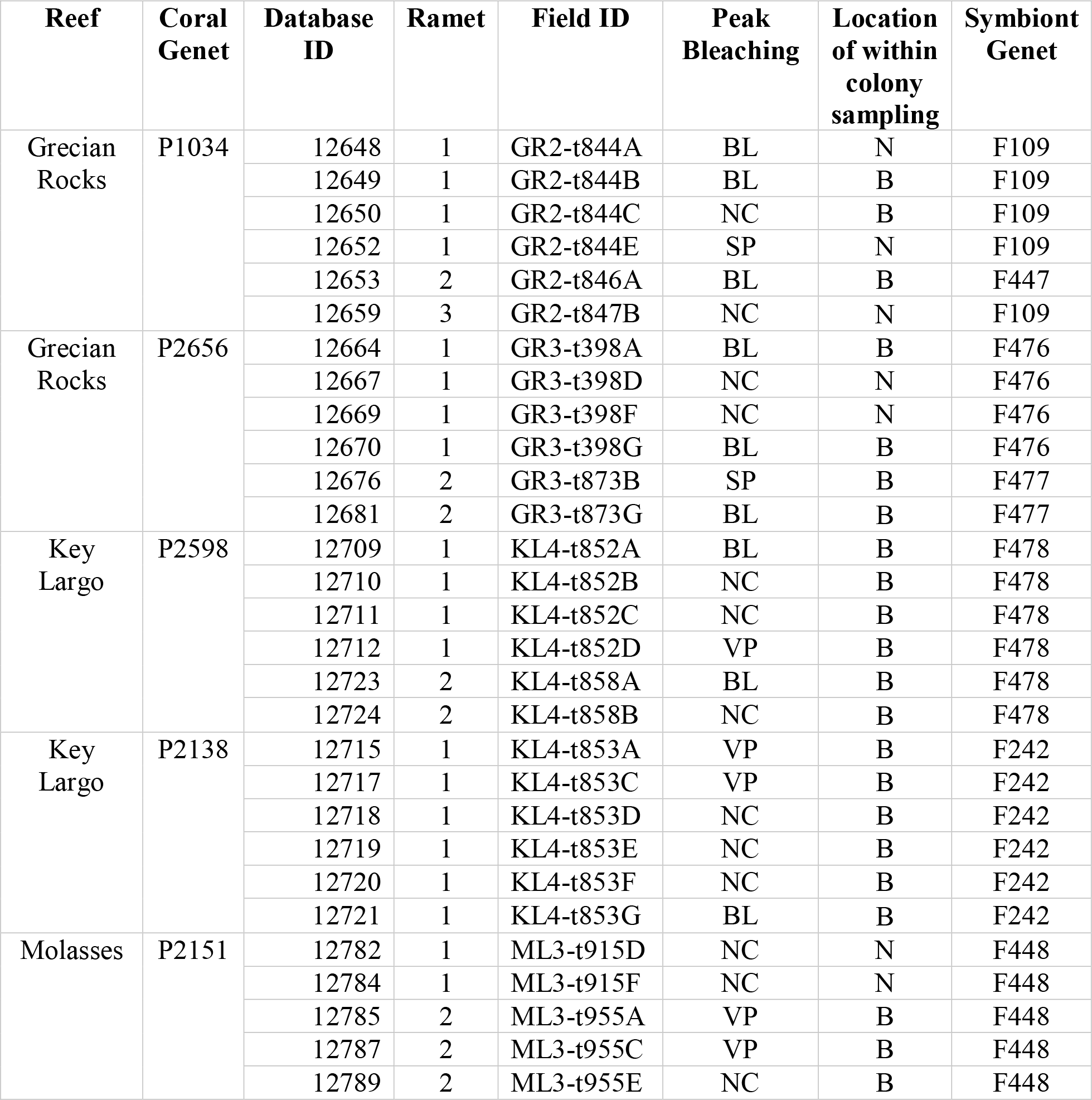
*Acropora palmata* colonies included in the MethylRAD analysis. Samples from 4 reefs in Florida contained 6 different coral genets and 8 *Symbiodinium fitti* genets. Phenotypic data included bleaching condition (BL=bleached, VP=very pale, NC=normal color, SP=slightly pale), and sampling location within a colony (Branch=B, not a branch=N).

| Reef | Coral Genet | Database ID | Ramet | Field ID | Peak Bleaching | Location of within colony sampling | Symbiont Genet |
| --- | --- | --- | --- | --- | --- | --- | --- |
| Grecian Rocks | P1034 | 12648 | 1 | GR2-t844A | BL | N | F109 |
|  |  | 12649 | 1 | GR2-t844B | BL | B | F109 |
|  |  | 12650 | 1 | GR2-t844C | NC | B | F109 |
|  |  | 12652 | 1 | GR2-t844E | SP | N | F109 |
|  |  | 12653 | 2 | GR2-t846A | BL | B | F447 |
|  |  | 12659 | 3 | GR2-t847B | NC | N | F109 |
| Grecian Rocks | P2656 | 12664 | 1 | GR3-t398A | BL | B | F476 |
|  |  | 12667 | 1 | GR3-t398D | NC | N | F476 |
|  |  | 12669 | 1 | GR3-t398F | NC | N | F476 |
|  |  | 12670 | 1 | GR3-t398G | BL | B | F476 |
|  |  | 12676 | 2 | GR3-t873B | SP | B | F477 |
|  |  | 12681 | 2 | GR3-t873G | BL | B | F477 |
| Key Largo | P2598 | 12709 | 1 | KL4-t852A | BL | B | F478 |
|  |  | 12710 | 1 | KL4-t852B | NC | B | F478 |
|  |  | 12711 | 1 | KL4-t852C | NC | B | F478 |
|  |  | 12712 | 1 | KL4-t852D | VP | B | F478 |
|  |  | 12723 | 2 | KL4-t858A | BL | B | F478 |
|  |  | 12724 | 2 | KL4-t858B | NC | B | F478 |
| Key Largo | P2138 | 12715 | 1 | KL4-t853A | VP | B | F242 |
|  |  | 12717 | 1 | KL4-t853C | VP | B | F242 |
|  |  | 12718 | 1 | KL4-t853D | NC | B | F242 |
|  |  | 12719 | 1 | KL4-t853E | NC | B | F242 |
|  |  | 12720 | 1 | KL4-t853F | NC | B | F242 |
|  |  | 12721 | 1 | KL4-t853G | BL | B | F242 |
| Molasses | P2151 | 12782 | 1 | ML3-t915D | NC | N | F448 |
|  |  | 12784 | 1 | ML3-t915F | NC | N | F448 |
|  |  | 12785 | 2 | ML3-t955A | VP | B | F448 |
|  |  | 12787 | 2 | ML3-t955C | VP | B | F448 |
|  |  | 12789 | 2 | ML3-t955E | NC | B | F448 |

| **Reef** | **Coral Genet** | **Database ID** | **Ramet** | **Field ID** | **Peak Bleaching** | **Location of within colony sampling** | **Symbiont Genet** |
| --- | --- | --- | --- | --- | --- | --- | --- |
|  |  | 12793 | 3 | ML3-x01C | BL | B | F448 |
| Sand Island | P1002 | 12808 | 1 | SI1-t959C | BL | B | F449 |
|  |  | 12810 | 1 | SI1-t959E | NC | B | F449 |
|  |  | 12811 | 1 | SI1-t959F | VP | N | F449 |
|  |  | 12812 | 1 | SI1-t959G | NC | N | F449 |
|  |  | 12813 | 1 | SI1-t959H | BL | N | F449 |
|  |  | 12814 | 1 | SI1-t959I | NC | B | F449 |

### *Symbiodinium* Microsatellite Analysis

*Symbiodinium ‘fitti’* DNA was amplified with primers found in Pinzón *et al.* (2011) (n = 10 loci, Table S1, Supporting information). Three additional loci for *S. ‘fitti’* were developed *de novo* (Table S1, Supporting information). Alleles were fluorescently visualized and sized with internal standards on a PRISM 3100 Genetic Analyzer (Applied Biosystems). Briefly, 25–50 ng of template DNA was added to Polymerase Chain Reactions (PCR) containing 1× Standard Taq Buffer (New England Biolabs), 2.5 mm MgCl (New England Biolabs), 0.5 mg/mL Bovine serum albumin (New England Biolabs) and 0.75 U of Taq (0.325 U for primers 31, 32 and 41; New England Biolabs). Primer concentrations of 200 nm of each primer were added to reactions involving loci A3Sym_01, 18, 27 and 28, whereas a primer concentration of 93 nm was added to reactions for primers 31, 32, 41. Primer concentrations of 50 nm of the tailed forward primer (see Table S1, Supporting information), 150 nm of the reverse primer and 75 nm of the dye labelled T-oligonucleotide were used for amplifying loci A3Sym_01, 02, 03, 07, 08, 09 and 48. All loci were amplified using the following thermal cycle profile: 94 °C for 2 min (1 cycle); 94 °C for 15 s, primer-specific annealing temperature (Table S1, Supporting information) for 15 s, 72 °C for 30 s (31 cycles); 72 °C for 30 min on a Mastercycler gradient thermal cycler (Eppendorf, Westbury, NY, USA). See Table S2 for microsatellite genotype allele calls.

### 16S Library Preparation and Bioinformatics

The hypervariable V1 and V2 regions of the 16S rRNA gene were amplified using a general bacterial primer pair with Illumina CS1 and CS2 Fluidigm adapters. [27F 5’-TCGTCGGCAGCGT CAGATGTGTATAAGAGACAGAGAGTTTGATCMTGGCTCAG-3’ and 355R 5’-GTCTCGTGGGCTCGGAGATGTGTATAAGAGACAGGCTGCCTCCCGTAGGAGT-3’] (Rodriguez-Lanetty et al. 2013). Polymerase chain reactions were performed using 2 U Gotaq (Promega), 1X Gotaq buffer (Promega), 0.25 mM dNTPs (Bioline), 2.5 mM MgCl2, and 0.25 μM of each primer. Initial denaturation was performed at 95°C for 5 min followed by 30 cycles of 95°C for 30 sec, 51°C for 1 min, and 72°C for 1 min. Final extension was performed at 72°C for 7 min. PCR products were checked on a 1% agarose gel for a band near 450bp. Libraries were prepared at the DNA services facility at the University of Illinois, Chicago and sequenced on a MISEQ2500 platform. Genets P1034, P2138, P1002, and P2151 were run on one chip, and Genets P2656 and P2598 were run together on the second Miseq chip.

Paired-end sample data with quality scores were imported and then reads were joined using vsearch join-pairs and quality filtered using quality-filter q-score-joined within Qiime2 (Bolyen *et al.* 2018; Caporaso *et al.* 2010). Chimeras were removed and sequences denoised using deblur with p-trim-length parameter of 230 which truncates the read at this position. Illumina runs were analyzed separately in Qiime2 and combined after the deblur analysis. The Naive Bayes classifier was trained on only the region of the target sequences that was sequenced and taxonomy assigned using Silva 132 reference sequences clustered at 99% sequence similarity. The output was a table containing operational taxonomic units (OTU). OTUs were normalized using the QIIME2 plugin, q2-perc-norm (Gibbons *et al.* 2018), a model-free approach that corrects batch effects in case-control microbiome studies, with condition of bleaching (Normal color/slightly pale (Not bleached) vs bleached/very pale (Bleached)) as the case-control. Correction for batch effects was necessary because genet was confounded with Miseq run.

Differences in microbiome community composition among genet, reef, or bleaching condition were tested using permutational multivariate analysis of variance (PERMANOVA). Bray–Curtis dissimilarity matrices were created using the vegdist() function (method = “bray”) and PERMANOVA tests (999 permutations) were done using the adonis() function in the vegan package (Dixon 2009). The R vignette variancePartition (Hoffman & Schadt 2016) was then used to fit a linear mixed model for each order/genus of bacteria and partition the total variance into the fraction attributable to each phenotype in the design, including genet, location of sampling (branch vs not branch), and the condition of the sample (bleached vs not bleached).

### Methylrad library preparation and Bioinformatics

Coral tissue samples were snap frozen in liquid nitrogen and kept at −80° C before extraction using the DNeasy Blood & Tissue Kit (QIAGEN) with the following modifications. Time of incubation in extraction buffer was increased to 16-20 hours. Further, only the second elution was retained for library production as this fraction contained the high molecular weight (HMW) DNA. A total of 100ng of HMW genomic DNA was digested with FspEI (NEB) at 37°C for 4 hours. Five μL of digested DNA was run on an acrylamide gel to verify digestion. Ligation was performed in a 30 μL reaction with 20 μL of digested DNA, 0.2 μM each of two adaptors, 1 mM ATP and 800 U of T4 DNA ligase (NEB). Ligation was carried out at 4° C overnight. Adaptor and primer sequences are provided in the supplementary material (Table S3). Ligation products were amplified in 40 μL reactions containing 15 μL of ligated product, 0.2 mM of each primer (p1 and p2), 0.3 mM dNTP, 1X Phusion HF buffer and 0.4 U Phusion high-fidelity DNA polymerase (NEB). PCR was performed in an Eppendorf Mastercycler with 22 cycles of 98°C for 5 s, 60°C for 20 s, 72°C for 10 s, and a final extension of 5 min at 72°C. The target DNA band (approx. 100 bp) was excised from a 10% TBE polyacrylamide gel, and the DNA was diffused from the gel in 50 μL of nuclease-free water for 12 h at 4°C. Barcodes were added in a second PCR amplification (total volume =20 μL), which contained 7 μL of gel-extracted PCR product, 0.2 mM of each primer (p3 and index primer), 0.3 mM dNTP, 1X Phusion HF buffer and 0.4 U Phusion high-fidelity DNA polymerase (NEB). Seven PCR cycles with the same profile outlined above were performed. PCR products were purified using QIAquick minElute PCR purification kit (Qiagen) and then with Ampure XP beads (Agencourt). Libraries were run on an Illumina HiSeq 2500 sequencer using 50 nt single read sequencing.

Raws reads were converted from Illumin 1.8+ to FastQSanger quality scores using FastQ Groomer (Blankenberg *et al.* 2010). Illumina adaptors were removed with Trim Galore! (The Babraham Institute by <u>@FelixKrueger</u>) and Trim (Galaxy Version 0.0.1) was used to clip 2 base pairs off both ends. A reference-based approach was used by extracting FspE1 methyl sites from an *Acropora palmata* genome assembly (version 1.0, (Kitchen *et al.* 2018)) using a custom Perl script (provided by Shi Wang) and reads were mapped against these reference sites using Bowtie2 (default settings, (Langmead & Salzberg 2012)). Samtools IdxStats (Li *et al.* 2009) was used to tabulate mapping stats into counts of mapped reads per reference sites for each sample.

Differential DNA methylation analysis was performed in EdgeR. Within the EdgeR vignette, data is converted to reads per million and filtered for only those methylation sites that have at least 10 reads per million in at least 6 samples. Then normalization factors were calculated to correct for the different compositions of the samples. The effective library sizes are then the product of the actual library sizes and these factors. After library size normalization the data was subjected to non-parametric batch normalization with the ComBat package for R software (http://www.bu.edu/jlab/wp-assets/ComBat/Abstract.html). Common, trended, and tagwise dispersions were estimated and negative binomial generalized linear models were fitted using both the likelihood ratio test and the more stringent quasi-likelihood (QL) F-test within EdgeR vignette. E-values were FDR corrected.

Methylated sites were matched against the *A. palmata* genome annotation. Methylated sites were analyzed with a Principal Components Analysis within the vegan package in R and visualized using ggbiplot (Wickham 2016). The R vignette variancePartition (Hoffman & Schadt 2016) was then used to fit a linear mixed model for each methylation site and partition the total variance into the fraction attributable to each phenotype in the design, including genet, location of sampling (branch vs not branch), and the condition of the sample (Bleached vs Not bleached) plus the residual variation.

Differentially methylated sites were tested for GO term functional enrichment with the R/Bioconductor package topGO (Rahnenfuhrer 2018) using the default “weight01” Alexa algorithm with the recommended cutoff of P < 0.05.

## Results

### Photographic Survey Results

Prior to the 2014 bleaching event, all sampled colonies had been monitored since at least 2010 and had not bleached until 2014. Following the 2014 bleaching seven of the 12 of the sampled ramets lost all or nearly all live tissue. Five ramets retained enough live tissue to evaluate their bleaching condition during the 2015 bleaching event. All five were observed to regain visually normal tissue pigmentation prior to late September 2015 when all five showed visual signs of bleaching, but generally less severe than in 2014. In many cases the pattern of bleached vs non-bleached areas on a colony was very similar to that observed in 2014 (Fig 2). Overall, visual surveys in 2015 showed that colonies responded similarly in 2014 and 2015. However, while in 2015, 10% (n=7) of colonies scored one or more classifications of less severely bleached than in 2014, and 54 % (n=37) of colonies (including samples that died) had greater bleaching classifications, indicating that many colonies were more severely impacted by the back-to-back bleaching of 2014-2015 (Table 3). Survey results thus provided little evidence of short-term acclimatization.

**Figure 2.**
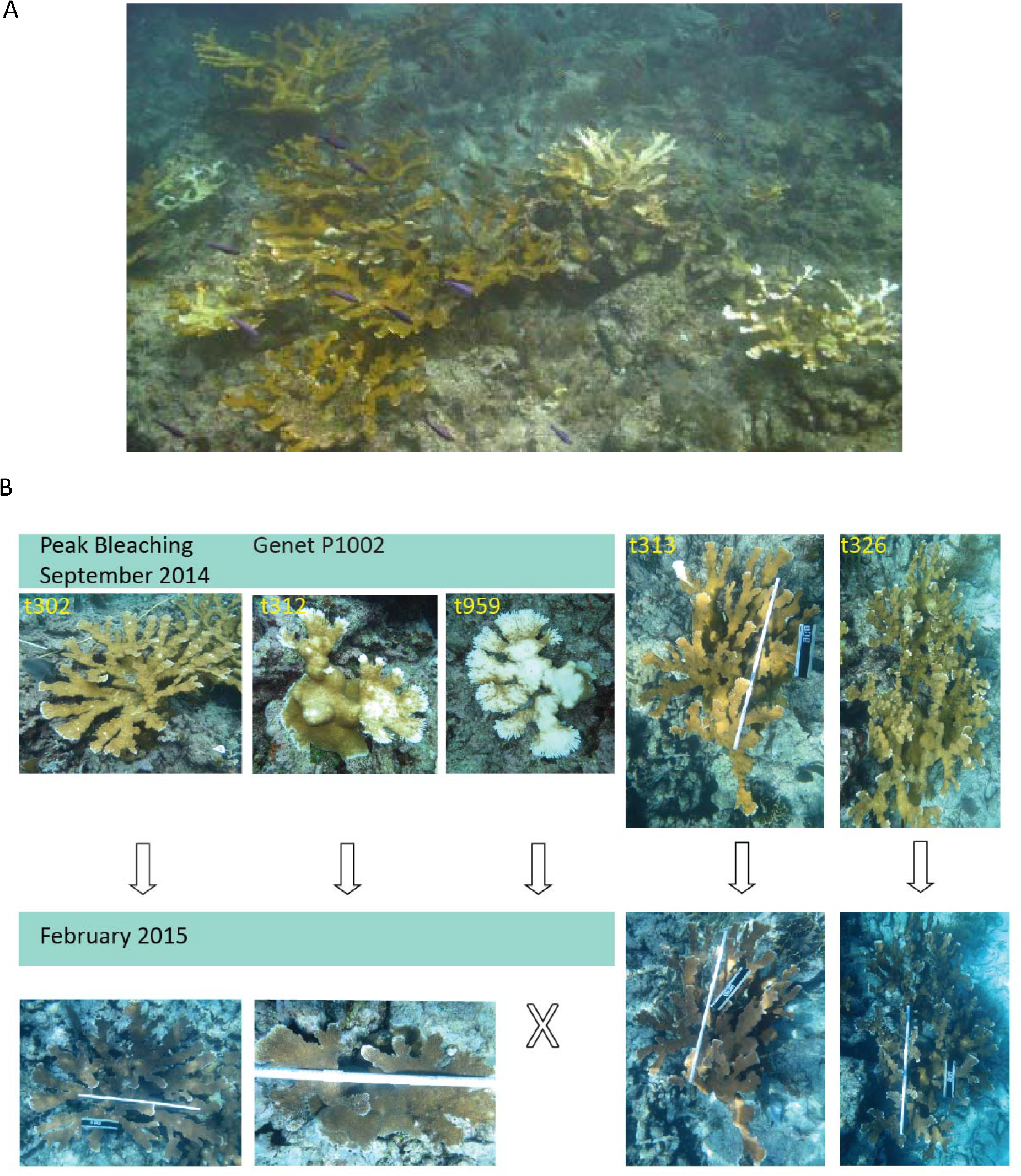
Heterogeneity in bleaching phenotype was observed among ramets (colonies) of the same *Acropora palmata* genet. A) Genet at Molasses Reef. B) Ramets of genet P1002 at Sand Island reef photographed during peak bleaching in September of 2014 and again in February 2015. Genet P1002 harbored a single strain of *S. ‘fitti* in all ramets. Images suggested that the heterogeneity in bleaching was due to micro-environmental differences that may have altered the underlying epigenome.

### *Symbiodinium* strain diversity was not correlated with bleaching response

Coral samples were genotyped at 13 microsatellite loci specific for *Symbiodinium ‘fitti’* (Pinzón et al. 2011) to determine whether *S. ‘fitti’* strain diversity was correlated with bleaching response. Each of the six samples taken per genet were dominated by one *S. ‘fitti’* strain with three exceptions (Table 1). Ramet 2 Branch A of Genet P1034 (Grecian Rocks) was dominated by *S. ‘fitti’* strain F447 rather than strain F109 like the other five samples from this genet. Ramet 2 Branches B and G of genet P2656 (Grecian Rocks) contained *S. ‘fitti’* strain F477 rather than strain F476 like the other 4 samples from this genet. These three within-genet *S. ‘fitti’* strain differences were not correlated with the peak bleaching event condition.

### Microbiomes differed among genets but only *Tepidiphilus* and *Endozoicomonas* abundances were correlated with peak bleaching response

Microbiomes differed significantly among genets at the taxonomic rank of order (Fig 3a) and genus (PERMANOVA analysis, Table 2). When blocking for the genetic background (ie Genet designation), microbiomes were not different between bleached and not bleached samples or between reefs at the taxonomic rank of order and genus (PERMANOVA analysis, Table 2). However, interestingly the microbiome did vary between sampling location within a colony, *i.e.* whether from branch or base locations (see below).

**Figure 3.**
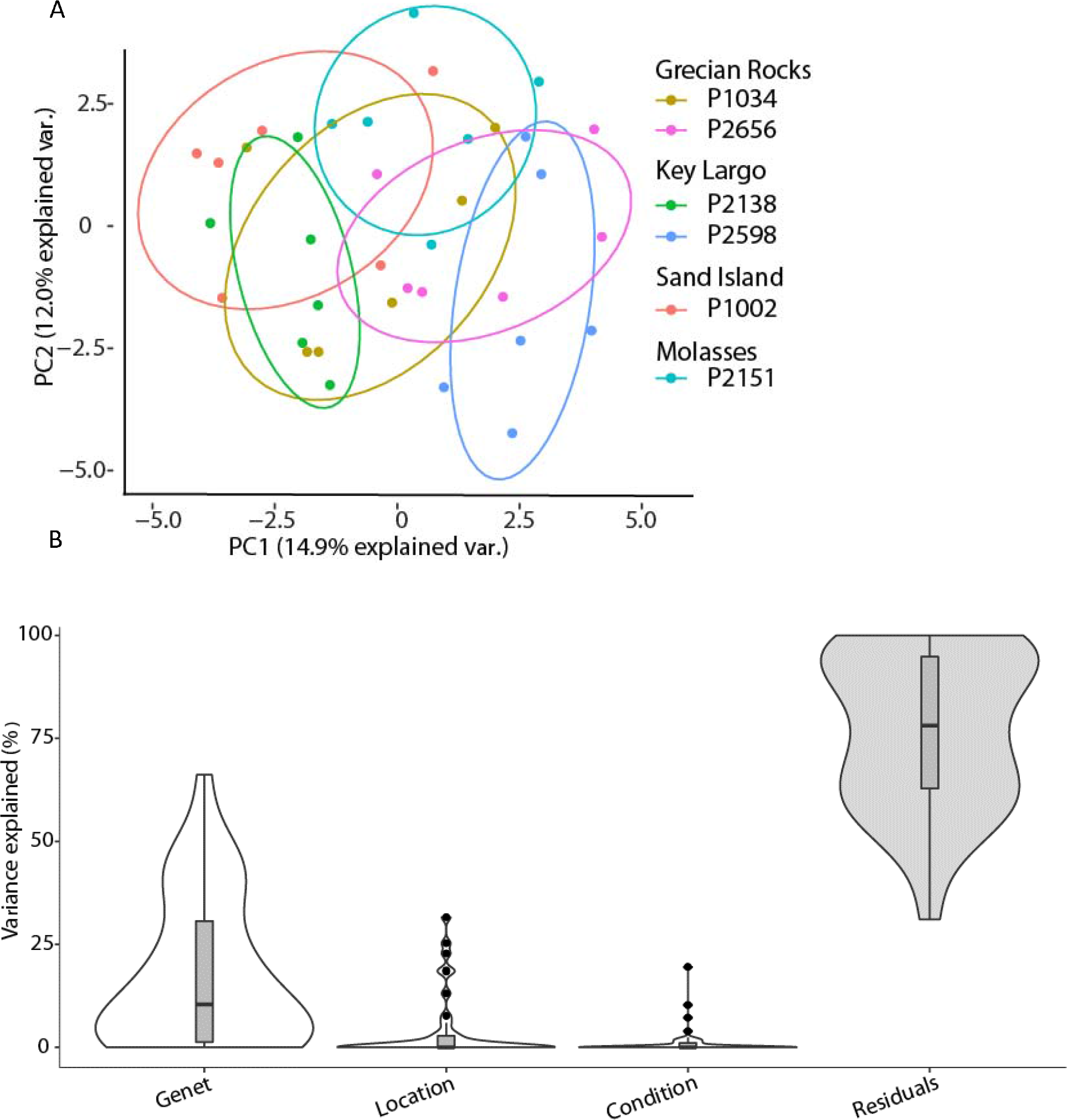
*Acropora palmata* genets are characterized by between and within genet variation in their microbiomes. A) Principal coordinate analysis of the normalized 16s tag sequence data generated using the package vegan and ggbiplot in R at the taxonomic rank of order. B) Violin plot showing the contribution of Genet, Location sampled (Branch, or not Branch), the Condition of sample (Bleached or not bleached) and the residuals to variation in the 16S tag normalized microbiome data at the taxonomic rank of order. Genet contributed most of the explained variation, followed by location and condition. Generated using the variancePartition package in R.

**Table 2.** PERMANOVA results based on Bray-Curtis dissimilarities using abundance data for microbiome community structure at the level of order (a) or genus (b). For the Reef, Sampling Location, and Bleaching condition comparisons data was blocked using Genet designation. Df-degrees of freedom; Sum Sq – sum of squares; Pseudo-F – F value by permutation; boldface indicates statistical significance with P<0.05, P-values based on 999 permutations.

| Factors | Df | Sum Sq | Pseudo-F | R2 | P |
| --- | --- | --- | --- | --- | --- |
| Genet | 5 | 0.69756 | 2.8573 | 0.3226 | **0.001** |
| Residuals | 30 | 1.46477 |  | 0.6774 |  |
| Total | 35 | 2.16232 |  | 1.0000 |  |
| Reef | 3 | 0.36385 | 2.158 | 0.16827 | 1 |
| Residuals | 32 | 1.79848 |  | 0.83173 |  |
| Total | 35 | 2.16232 |  | 1.00000 |  |
| Location | 1 | 0.14355 | 2.4177 | 0.06639 | **0.005** |
| Residuals | 34 | 2.01877 |  | 0.93361 |  |
| Total | 35 | 2.16232 |  | 1.00000 |  |
| Condition | 1 | 0.07035 | 1.1434 | 0.03254 | 0.125 |
| Residuals | 34 | 2.09197 |  | 0.96746 |  |
| Total | 35 | 2.16232 |  | 1.00000 |  |

B)
| Factors | Df | Sum Sq | Pseudo-F | R2 | P |
| --- | --- | --- | --- | --- | --- |
| Genet | 5 | 0.70612 | 3.0791 | 0.33914 | **0.001** |
| Residuals | 30 | 1.37596 |  | 0.66086 |  |
| Total | 35 | 2.08208 |  | 1.00000 |  |
| Reef | 3 | 0.3812 | 2.3906 | 0.18309 | 1 |
| Residuals | 32 | 1.7009 |  | 0.81691 |  |
| Total | 35 | 2.0821 |  | 1.00000 |  |
| Location | 1 | 0.11408 | 1.9709 | 0.05479 | **0.024** |
| Residuals | 34 | 1.96800 |  | 0.94521 |  |
| Total | 35 | 2.08208 |  | 1.00000 |  |
| Condition | 1 | 0.05864 | 0.9854 | 0.02816 | 0.169 |
| Residuals | 34 | 2.02344 |  | 0.97184 |  |
| Total | 35 | 2.08208 |  | 1.00000 |  |

Next, we determined if any of the variation was attributable to different phenotypic characteristics. Variables included polyp sampling location (branch vs not branch), sample peak bleaching condition (bleached vs. not bleached), and genetic background (genet). The largest partition, not including residuals, was to genetic background followed by within-colony sampling location, (branch vs not branch), and then bleaching condition, at both the taxonomic rank of order and genus (Fig 3b). The top three bacterial genera that had the highest variation attributable to the host genotype were Methylobacterium (σ^2^=60%), Rhodobacteraceae (σ^2^=52%), and Alteromonas (σ^2^=48.7%). The top three bacterial genera that had the highest variation attributable to the location of the samples (branch vs. not branch) were Cyclobacteriaceae (σ^2^=41.6%), Tepidiphilus (σ^2^=25.9%), and Amoebophilus (σ^2^=23.1%). There were only two bacterial genera in which variance attributable to the bleaching condition of the coral exceeded 5%; this included *Tepidiphilus* (σ^2^=12.1%) and *Endozoicomonas* (σ^2^=6.8%). Examining the sub OTU sequence output from Qiime2 focusing on *Tepidiphilus* and *Endozoicomonas* revealed only one strain for both.

### The methylome differs among coral host genets

Image analyses of bleached and unbleached portions of the colonies suggested that micro-environmental differences such as shading or exposure to water movement might have contributed to differential bleaching responses (Figure 2). Thus, we investigated an epigenetic mechanism of acclimatization, DNA methylation, to address whether different portions of a colony or genet had acclimatized to micro-environmental differences.

A total of 28,797 sites were analyzed for differential methylation, and 64.4% (n=18,538) of these sites are within predicted genes in the *A. palmata* genome (version 1). Variation attributable to different phenotypic characteristics was determined, and the largest partition, not including residuals, was to host genetic background followed by within-colony polyp sampling location, ie whether a branch location or not (Fig 4b), and lastly whether the polyps were previously bleached or not bleached. To determine the biological function of genes in which methylation was more abundant, a GO-enrichment analysis was completed comparing all methylsites (n=6,848 had GO term annotations out of 28,797 sites) to all annotated genes in the *A. palmata* genome (n=25,102 had GO term annotation). Significant biological functions included RNA binding, zinc ion binding, transition metal binding, sequence-specific DNA binding, and oxidoreductase activity (Table S4).

Genets showed large and consistent differences in methylation patterns and clustering analyses clearly grouped samples belonging to the same coral genet together, but also showed some variation of methylation within genets (Fig 4a). On average 9.5% (n=2741, QL F-test) of sites were significant for differential methylation between genets and of these sites, 57.8% were within predicted genes (n=1584). In a GO enrichment of the differentially methylated sites between genets, the top 5 biological processes were the negative regulation of molecular function, the negative regulation of hydrolase activity, the modification by symbiont of host morphology or physiology, the negative regulation of proteolysis, and the regulation of DNA-binding transcription factor activity (Top 20 nodes, Biological processes: Supplemental Table 5, Molecular Function: Supplemental Table 6).

**Figure 4.**
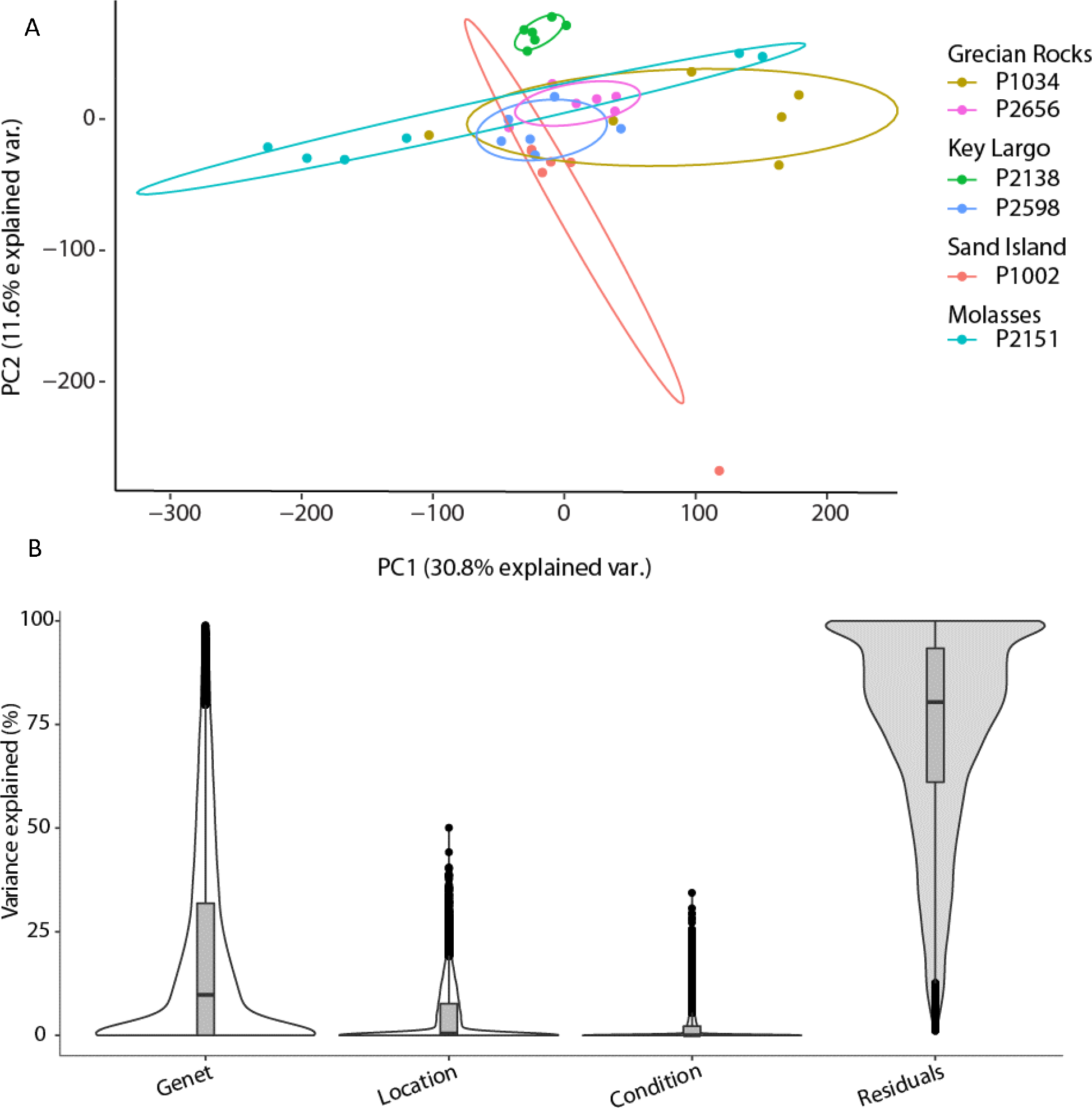
*Acropora palmata* genets are characterized by between and within genet variation in genome methylation. A) Principal coordinate analysis of the normalized coral methylome data generated using the package vegan and ggbiplot in R. B) Violin plot showing the contribution of Genet, Location sampled (Branch, or not Branch), the Condition of sample (bleached or not bleached) and the residuals to variation in the normalized coral methylome data. Genet contributed most of the explained variation, followed by location and condition. Generated using the variancePartition package in R.

### Reef, peak bleaching, and polyp location within colonies each explained some of the variation in differential methylation among host genets

Next, we determined whether there were methylation changes attributable to the reef environment. On average 5% (n=1511, QL F-test) of sites were differentially methylated among reefs. Unfortunately, we could not block for genetic background in the QL F-test analysis because not all reefs were represented by multiple genets. However, a venn diagram analysis comparing which CCGG sites are differentially methylated between reefs versus those differentially methylated between genets, showed there was a 40.1% (n=1218) overlap between the factors. Only 9.7% (n=293) of the differentially methylated sites were exclusively found in the reef comparison compared to 50.2% (n=1523) were found exclusively in the genet comparison (Supplemental Figure 2). A GO enrichment for biological processes for these differentially methylated sites exclusively found in comparison between reefs but not between genets (n=293), showed that the top 5 processes were in autophagy and the regulation thereof, amine transport including both regulation and positive regulation, and the regulation of cellular catabolic processes. (Top 20 nodes, Biological processes: Table S7, Molecular Function: Table S8).

We then examined whether there are conserved methylation differences shared among all genets that were attributable to whether an area of a colony previously bleached or did not bleach. In this comparison, we included 18 bleached and 18 non-bleached samples and blocked for genet. One methylation site (Segkk362_pilon-108100) was significantly different (p<0.01, FDR corrected) when using a likelihood ratio test, but not when using the more stringent quasi-likelihood (QL) F-test. This methylation site lies within a genomic region that does not have a gene prediction or annotation in *A. palmata*. The closest gene annotation, at a little over 10,000 bp away, is Transposon TX1 uncharacterized 149 kDa protein as found in the scleractinian coral, *Stylophora pistillata*.

Next, we looked at whether samples collected from branches versus those collected from the base or trunk of the colony (i.e. not branch) had conserved methylation differences shared among all genets. In this comparison (n= 26 branch, n= 10 not branch) when blocking for genet, there was one methylation site (Segkk1116_pilon-156192) that was differently methylated (p<0.001, FDR corrected) when using the more stringent quasi-likelihood (QL) F-test and 84 methylation sites when using the likelihood ratio test (Table S9). Segkk1116_pilon-156192 is within the gene Galactosylceramide sulfotransferase. Of the 84 differentially methylated sites found in the likelihood ratio test, 61.9% (n=52) are within predicted genes. A GO-enrichment analysis of these differentially methylated sites, using TopGO in R, revealed a significant enrichment for the biological processes (Supplemental Table 10) of AMP biosynthetic and metabolic processes, regulation of fibroblast apoptotic processes, excretion, protein-chromophore linkage, the regulation of viral genome replication, viral life cycle, and viral processes, the regulation of alternative nMRNA splicing via spliceosome, immune effector processes, the negative regulation of apoptotic processes and programmed cell death (for enrichment in Molecular Function see Table S11).

## Discussion

Evolutionary rescue of populations depends on their ability to produce phenotypic variation that is heritable and adaptive. DNA mutations are the best understood mechanisms to create phenotypic variation, but other, less well-studied mechanisms exist. Here we, for the first time, applied a new technique to assay methylation status in wild corals and have found that coral genets show large and consistent differences in the way they methylate their DNA even when growing on the same reef, suggesting that methylation patterns are inherited (Liew *et al.* 2018b). Further, there was significant within colony variation in methylation state that was correlated with the location of the polyps within colonies and the bleaching response of the colonies. Remarkably, there was some correspondence between genes previously shown to be differentially expressed, and those that were differentially methylated indicating that methylation difference may translate into gene expression differences. These results are novel because they reveal a pathway by which these long-lived corals can modify their phenotypes in response to the environment. Because these corals can produce large monoclonal stands, these modifications produce mosaics of phenotypes despite low genotypic diversity.

Environmental conditions frequently change over the live span of reef-building coral genets. Phenotypic plasticity is a common trait in corals; however, the mechanisms by which corals achieve this plasticity are not well understood. This can be partly attributed to the difficulty of separating the host response from those of algal symbionts and other members of the microbiome. *A. palmata* colonies had variable responses to increased water temperatures yet, the identity of the associated symbiont strain did not explain this plasticity. In addition, photographic survey results showed little evidence of short-term acclimatization. Interestingly, the abundance of two species of prokaryotes, *Tepidiphilus* and *Endozoicomonas* differed among samples depending on their bleaching response and this supports findings that *Endozoicomonas* plays an important role in the functioning of the coral holobiont (Pollock *et al.* 2018).

### Microbial assemblages and symbiotic dinoflagellates are minimally correlated with bleaching condition

Corals live in an intimate symbiosis with algae in the family Symbiodiniaceae. While some coral species can host several species of algae, the elkhorn coral, *A. palmata* usually hosts *S. ‘fitti’* and furthermore ramets of the same coral genet often retain the same symbiont strain (Baums *et al.* 2014). This was also the case here. Genets differed in the *S. ‘fitti’* strain they hosted but within each genet, there was remarkable homogeneity. These findings allow us to discount genetic diversity of *S. ‘fitti’*, at least at the strain level, as an explanation for differential bleaching response. However, variable densities of symbiotic dinoflagellates within the host tissues may have influenced bleaching susceptibility (Kemp *et al.* 2014; Stimson *et al.* 2002). We do not have pre-bleaching measurements of symbiont density in tissues, but this may have contributed to variable bleaching within a colony.

Prokaryotic members of the coral microbiome play important roles in coral nutrition, element cycling, and disease responses (Peixoto *et al.* 2017). We thus speculated that the differential bleaching response within and between colonies might be attributable to differences in the microbiome species composition. However, there were only two bacterial genera in which variance attributable to the bleaching condition of the coral exceeded 5%: *Tepidiphilus* (σ^2^=12.1%) and the coral symbiont *Endozoicomonas* (σ^2^=6.8%). Each taxon was represented by one strain only. There are three proposed functions of *Endozoicomonas* in the host coral, including the maintaining the structure of the host microbiome, nutrient acquisition and provision, and a role in host health and/or disease (Neave *et al.* 2017). Evidence for the role in coral health was demonstrated through the altered abundances of *Endozoicomonas* in multiple scleractininan corals in relation to seawater pH (Webster *et al.* 2016), sedimentation and wastewater runoff (Ziegler *et al.* 2016), and occurrence of diseased lesions (Meyer *et al.* 2014). Differences in the microbiome that predated or immediately followed the bleaching event were obscured, because we were only able to sample tissues once they had recovered from bleaching. Previous studies looking at stress events in corals found a shift of the coral holobiont to a more potentially pathogenic state with disease associated bacteria and fungi (Thurber *et al.* 2009), including the fungi *Ascomycota* (Thurber *et al.* 2009) and the bacteria taxa vibrionales (Thurber *et al.* 2009; Zaneveld *et al.* 2016) and oscillatoriales (Zaneveld et al. 2016). A shift to a pathogenic state was not observed in our colonies indicating that whatever impacts bleaching may have had on the microbiome, those effects were difficult to detect six weeks post-stress using a standard 16S microbiome analysis approach.

### Variation in methylation patterns by Genet

*Acropora palmata* genets differ greatly in their genome methylation patterns even when living on the same reef (Fig 2). This suggests that methylation patterns were inherited, otherwise a shared environment post-fertilization should lead to shared methylation patterns among genets. Evidence for inherited germline methylation patterns in corals is accumulating. Widespread depletion of CpG dinucleotides was observed in *Acropora millepora* and is a signature for historic germline DNA methylation. (Dixon *et al.* 2014). A recent study in *Platygyra daedalea* for the first time demonstrated intergenerational inheritance of DNA methylation patterns in corals, from parent to sperm and evidence for maternal and paternal effects in larvae from reciprocal crosses (Liew *et al.* 2018a).

Symbiont-host interactions may also influence host genome methylation patterns. Even genets that grew near each other on the same reef hosted a different strain of *S. ‘fitti’* while ramets of the same genet usually shared an *S. ‘fitti* strain. Although previously undocumented, *S. ‘fitti’* strains may differentially alter the host methylome. Supporting evidence for this comes from the genet by genet comparison gene ontology enrichment analysis which showed that the category “modification by symbiont of host morphology or physiology and modulation by symbiont of host cellular processes” was enriched. Differentially methylated genes in this category included the Homeodomain-interaction protein kinase 2, eIF-2-alpha kinase GCN2 (3 separate methylation sites within this gene), Gag-Pol polyprotein (2 separate methylation sites within this gene), and TNFAIP3-interating protein 1. Genotype/genotype interactions between Symbiodiniaceae strains and host genets and their effects on host methylomes deserve further study (reviewed in Parkinson & Baums 2014)

### Methylation patterns vary between locations within the colony

Some of the variance in genome methylation within genets was attributable to long-term microenvironmental conditions between polyp locations within a colony rather than eukaryotic or bacterial symbiont community composition. Complex skeletal morphologies and varying tissue layer thickness create a variety of intracolonial light microniches, but in general the top of a branch will experience significantly higher solar irradiance than the base or trunk of a colony (Kaniewska *et al.* 2011; Wangpraseurt *et al.* 2012; Warner & Berry-Lowe 2006) and therefore polyps on the tops of branches have an increased need to avoid the damaging effects of excess light energy and the resulting oxidative stress. In *A. globiceps*, *Symbiodinium* densities were consistent between internal and external branches but varied with depths (greater densities at lower depths) (Ladrière et al. 2013). The host coral can regulate symbiont densities through nutrient limitation or through digesting or expelling the excess symbionts to maintain low and consistent densities (Dunn *et al.* 2002; Falkowski 1993; Muscatine *et al.* 1998) as one option to reduce oxidative stress in areas or times of higher light exposure (Fitt *et al.* 2000).

Abnormally warm temperatures accompanied with high irradiance can cause a breakdown in the coral-symbiotic algae symbiosis resulting in the expulsion of the algae, a process referred to as bleaching. We observed a higher incidence of bleaching in samples from branch regions (58%) versus those collected from the base (30%) in this study. In addition, there is also strong evidence for a division of labor between coral branch tips and bases in their gene expression (Hemond *et al.* 2014). Interestingly, Galactosylceramide sulfotransferase was significantly differentially methylated between polyps sampled from branches versus other locations within colonies such as base or trunk. This result was obtained with both LRT and QLF test statistics. Galactosylceramide sulfotransferase is involved in sphingolipid metabolism and was also differentially expressed by colony position, being upregulated in branch tips in *Acropora palmata* and *A. cervicornis* (Hemond *et al.* 2014). Sphingolipid metabolism may be involved with the regulation of algal symbionts. In anemones, the sphingosine rheostat can regulate the balance between stability and dysfunction in the cnidarian-dinoflagellate partnership (Detournay & Weis 2011).

Symbiodiniaceae in shallow corals must dissipate four times more light energy than what is needed for photosynthesis on a bright summer day (Gorbunov *et al.* 2001). This excess light energy absorbed by chlorophyll can be dissipated through heat loss, reemitted as fluorescence, or decayed via the chlorophyll triplet state that produces reactive oxygen species as a byproduct. Here, we identified differentially methylated sites by polyp location that were overrepresented in the gene ontology categories of AMP biosynthetic and metabolic processes. The site is in the gene Adenylosuccinate lyase which catalyzes two key steps in AMP synthesis. cAMP induces gene transcription through the activation of cAMP-dependent protein kinase (PKA) and subsequently the activation of transcription factors including CREB (cAMP response element binding proteins)/ATF transcription factor family members such as CREM and ATF1 by phosphorylation by PKA. In a differential gene expression analysis of *A. palmata* fragments kept in complete darkness for 9 days compared to controls, two annotated genes were identified, cAMP-responsive element modulator and cyclic AMP-dependent transcription factor ATF-4 (DeSalvo *et al.* 2012). The ATF-4 transcription factor responds to oxidative stress and amino acid starvation (Harding *et al.* 2003).

The coral skeleton serves as an efficient light capturing devise and colony and polyp morphology determines the light levels experienced by the intra-cellular symbionts (Swain *et al.* 2018). Symbiodiniaceae can also maximize light absorption and utilization by increasing photosynthetic pigments and photosynthetic efficiency in low-light acclimated corals (Falkowski & Dubinsky 1981). Among the enriched gene ontology terms between branch and non-branch polyp locations was the category of Protein-chromophore linkage. The differentially methylated gene was in cyrptochrome-1. Cryptochromes are flavoproteins that are sensitive to blue light. They regulate the circadian clock in plants and animals. Eight core circadian genes have been identified: Casein kinase 1e (CK1e), Cryptochrome1 (Cry1), Cryptochrome2 (Cry2), Period1 (Per1), Period2 (Per2), Period3 (Per3), Clock, and BMAL1 (brain and muscle ARNT-like protein, Arntl, MOP3). Cryptochrome 1 and 2 have been previously reported to display diurnal patterns of transcription in corals, with higher expression found in the light phase than in the dark (Hoadley *et al.* 2011; Levy *et al.* 2007; Levy *et al.* 2011). Cry1 in *A. millepora* is not under control of an endogenous clock whereas Cry2 is. Both have higher expression during the day (Brady *et al.* 2011). Circadian clock genes affect a large number of downstream processes and thus serve as important nodes in transcriptional networks (Dunlap 1999) and in the regulation of post-translational modifications (Gallego & Virshup 2007; Staiger & Koster 2011). Differential methylation of these genes may thus be an effective means to alter the transcription of a number of downstream pathways in response to differential light levels within colonies. Future research related transcription and differential methylation of these circadian clock genes in corals is warranted.

We unexpectedly found that the gene ontology terms for the regulation of viral genome replication, viral life cycle, and viral processes were enriched in the comparison between branch and non-branch locations. We are not aware of any data indicating that viral load differs within colonies. Because this GO term was enriched across genets and reefs, we would expect viral loads to differ systematically between branch and non-branch locations and this hypothesis deserves future testing. Interestingly, the bacterial communities did differ between the tips and the bases of colonies suggesting that the branches may harbor a specialized microbiome. There is contradicting prior evidence with respect to within colony variation of the prokaryotic community in corals. In a previous study on *A. palmata*, no detectable community-level difference were found among the prokaryotic microbiota of the uppermost, underside, and base of *A. palmata* (R^2^ = 0.20, p = 0.51) (Kemp *et al.* 2015). In contrast, considerable within colony variation of bacterial assemblages were found in *O. annularis* between the tops and the sides (Daniels *et al.* 2011). *O. annularis* also harbors a several species of Symbiodiniaceae, hence further research is required to understand what factors drive within coral-colony diversity of the prokaryotic community.

### Methylation patterns vary with bleaching history

Methylation variation within genets was also attributable, to some extent, to whether tissues had recently bleached. Bleaching itself was likely the result of the interaction of the acute heat stress and microenvironmental variation. The one significant methylation site was located within a genomic region that does not have a gene prediction or annotation. The closest gene annotation, at a little over 10,000 bp away, is Transposon TX1 uncharacterized 149 kDa protein as found in *Stylophora pistillata*. By the alteration of splicing and polyadenylation patterns or through functioning as enhancers or promoters, transposable element can exercise control over neighboring genes (Girard & Freeling 1999). Transposable elements are significantly differentially expressed in response to heat stress in corals (DeSalvo *et al.* 2010; Traylor-Knowles *et al.* 2017) and in plants (Ito *et al.* 2011; Pecinka *et al.* 2010). Overall, less of the variation in methylation was explained by previous bleaching suggesting that methylation changes may be effective in changing coral transcription in response to longer term changes in the environment rather than more acute stressors. Future research is required to correlate gene expression and methylation over a range of stress exposures and stress severity.

High fragmentation rates and acute stress events necessitate that *A. palmata* polyps acclimate to changes in environmental conditions. Our data suggest that acclimatization is partially achieved via differential methylation. We suggest here that differential genome methylation may be one of the mechanisms by which corals achieve their remarkable phenotypic plasticity in their natural environment. In a transplant experiment in *A. millepora* gene body methylation changed subtly, but much less than transcription. They also found strong associations between GBM and fitness, however GBM was not directly correlated to transcription resulting in the authors questioning what mechanism connects GBM to phenotype and fitness (Dixon *et al.* 2017). Yet, methylation is known to affect transcription factor binding both negatively and positively, and thus alter transcriptional regulation bi-directionally, making correlation to gene expression complicated (Yin *et al.* 2017).

A significant amount of the intra-genet variation in phenotypic stress response observed here remains to be explained. Ultra-deep sequencing of host and symbionts may reveal occurrence of somatic mutations correlated with the bleaching phenotype (Van Oppen *et al.* 2011) but non-mutation based mechanisms other than DNA methylation changes may also play a role (Goldsmith & Tawfik 2009; Payne & Wagner 2019). Future work on this and other marine foundation species may significantly advance our understanding of these mechanism in determining evolvability of threatened species.

## Acknowledgements

This work was supported by a grant from the National Science Foundation OCE-1516763to IB and DW and OCE-1537959 to IB. Coral collections were conducted under sampling permit number: FKNMS-2014-148-A2 and survey permit: FKNMS-2010-130-A1 from the Florida Keys National Marine Sanctuary. Thanks to Shi Wang for help with adaptation of the MethylRad protocol for corals and providing a custom Perl script for data analysis.

## Data accessibility

Short read sequencing data is pending availability at GEO NCBI.

## Author contributions

IB, MD and DW designed the study and obtained funding. DK, MD and DW performed research and analyzed the data. MD and IB wrote the paper. All authors edited the paper.

